# Herpes Simplex Virus type 1 Inflammasome Activation in Human Macrophages is Dependent on NLRP3, ASC, and Caspase-1

**DOI:** 10.1101/796235

**Authors:** Andrew H. Karaba, Alexis Figueroa, Guido Massaccesi, Sara Botto, Victor R. DeFilippis, Andrea L. Cox

## Abstract

The pro-inflammatory cytokines interleukin (IL)-1β and IL-18 are products of activation of the inflammasome, an innate sensing system, and important in the pathogenesis of herpes simplex type 1 (HSV-1). The release of IL-18 and IL-1β from monocytes/macrophages is critical for protection from HSV-1 based on animal models of encephalitis and genital infection, yet if and how HSV-1 activates inflammasomes in human macrophages is unknown. To investigate this, we utilized both primary human monocyte derived macrophages and human monocytic cell lines (THP-1 cells) with various inflammasome components knocked-out. We found that HSV-1 activates inflammasome signaling in pro-inflammatory primary human macrophages. Additionally, HSV-1 inflammasome activation is dependent on nucleotide-binding domain and leucine-rich repeat-containing receptor 3 (NLRP3), apoptosis-associated speck-like molecule containing a caspase recruitment domain (ASC), and caspase-1, but not on absent in melanoma 2 (AIM2), or gamma interferon-inducible protein 16 (IFI16). Ultraviolet irradiation of HSV-1 enhanced inflammasome activation, demonstrating that viral replication suppresses inflammasome activation. These results confirm that HSV-1 is capable of activating the inflammasome in human macrophages through an NLRP3 dependent process and that the virus has an NLRP3 specific mechanism to inhibit inflammasome activation in monocytes and macrophages.

**Author Summary:** The inflammasome is a multi-protein complex that forms in response to pathogens and cellular damage. Active inflammasomes recruit pro-caspase-1 via ASC and cleave the cytokine precursors pro-IL-1β and pro-IL-18 into mature IL-1β and IL-18. These cytokines serve to activate other immune cells that either repair the damage or attempt to clear the invading pathogen. Upon activation, the inflammasome also promotes an inflammatory form of cell death called pyroptosis. Herpes simplex virus type 1 (HSV-1) is a common human pathogen that can cause cold sores, genital ulcers, encephalitis, and blindness. HSV-1 infection leads to induction of IL-1β and IL-18, but whether it is capable of activating inflammasomes in macrophages, which play a role in severe forms of HSV-1 infection, was unclear. Here, we infected both primary human macrophages and a macrophage/monocytic cell line, THP-1 cells, with HSV-1. We found that HSV-1 does activate inflammasome signaling in macrophages in a process dependent on NLRP3, ASC, and caspase-1. This is important because it illustrates the mechanism by which HSV-1 infection leads to inflammasome activation in macrophages, known to be crucial for protection from severe disease in mouse models.

## Introduction

The ability to quickly recognize and respond to pathogens is essential to host survival. The first opportunity to do so lies in the innate immune response. One of the most essential aspects of this response is the recognition of pathogen associated molecular patterns (PAMPs) on the invading pathogen by pattern recognition receptors (PRRs) within host cells [1]. This interaction leads to a number of molecular and cellular signals that serve to protect the host on cellular and organism levels. One such innate signaling system is the formation of inflammasomes which are intracellular multi-protein complexes that regulate an inflammatory type of cell death called pyroptosis, as well as, the production of mature forms of the inflammatory cytokines IL-1β and IL-18 [2]. Macrophages and myeloid dendritic cells (mDCs) are the primary producers of these potent pro-inflammatory cytokines, which drive type 1 immunity in natural killer cells and T-cells [3]. The production of these cytokines requires two steps. The first step, sometimes referred to as priming, requires activation of the nuclear factor κB (NF-κB) pathway through the recognition of a PAMP leading to synthesis of components of the inflammasome, including pro-IL-1β, pro-IL-18, and pro-caspase-1. The second step involves PRR activation, oligomerization, and assembly of the inflammasome. This takes place through one of multiple receptor or adapter proteins that recognize various PAMPs or danger-associated molecular patterns (DAMP). These include members of the nucleotide-binding domain and leucine-rich repeat-containing receptors (NLR) family of proteins, absent in melanoma 2 (AIM2), and pyrin. NLRP3 responds to a diverse group of PAMPs and DAMPs, particularly viral RNA [4–7]. In contrast, AIM2 is activated after binding to cytoplasmic double stranded DNA (dsDNA) [8]. Recognition of an appropriate PAMP or DAMP by one of these adapter proteins leads to apoptosis-associated speck-like molecule containing a caspase recruitment domain (ASC) assembly and oligomerization followed by pro-caspase-1 recruitment to the complex. Pro-caspase-1 autocatalyzation to active caspase-1 allows for cleavage of pro-IL-1β and pro-IL-18 to their active forms, IL-1β and IL-18, and then secretion into the extracellular space (reviewed in [2,9,10]). There are multiple sensors and caspases that can lead to inflammasome cytokine release, but the caspase-1 pathway is thought to be the most relevant in viral infection [11,12].

A number of viruses are known to activate the inflammasome, including influenza, hepatitis C, HIV, and herpesviruses [10,13]. Herpes simplex virus type 1 (HSV-1) is a neurotropic alphaherpesvirus that predominantly infects epithelial cells and neurons, but has broad cell tropism [14]. Specifically, it can infect macrophages, which are one of the predominant cell types that infiltrate the eye after corneal infection and are a critical cell in the innate immune response to HSV-1 and other viruses [15–17]. Furthermore, monocytes/macrophage production of IL-1β and IL-18 is critical to prevent severe HSV disease in encephalitis, keratitis, and vaginal infection in mouse models [17–19]. Therefore, understanding how HSV-1 activates the inflammasome in these cells is key to developing a comprehensive view of HSV-1 pathogenesis. A previous study demonstrated that the HSV-1 viral tegument protein VP22 specifically blocks AIM2 inflammasome activation and signaling in the THP-1 monocyte/macrophage cell line despite production of IL-1β, leaving the mechanism of HSV-1 activation of the inflammasome in these cells to be defined [20,21]. Thus, it remains unclear which adapters are required for HSV-1 induction of inflammasome activation in macrophages.

Here, we report that HSV-1 activates inflammasome signaling, as measured by IL-18, in both primary human macrophages and THP-1 cells. Additionally, this activation requires NLRP3, ASC, and Caspase-1, but not AIM2 or IFI16.

## Results

### HSV-1 activates the inflammasome in primary human monocyte derived macrophages

HSV-1 is known to activate the inflammasome in THP-1 cells, but the mechanisms is unknown and it has not been studied extensively in primary human macrophages [20,21]. To determine if HSV-1 is capable of activating the inflammasome in primary human monocyte derived macrophages (MDM), we inoculated both unstimulated macrophages (referred to as M0) and macrophages incubated for 24 hours with IFNγ (referred to as M1) [22,23] with HSV-1, media alone, or nigericin and LPS (a potent activator of the NLRP3 inflammasome [24]) and then measured IL-18 in supernatants 24 hours later. The M0 MDMs did not produce significant amounts of IL-18 after HSV-1 infection (**Fig. 1a**), but did in response to nigericin and LPS stimulation (**Supp Fig. 1**). However, IL-18 was detected after HSV-1 infection in the M1 MDMs (**Fig. 1b**). Additionally, in response to nigericin and LPS, M1s produced more IL-18 than M0s suggesting that IFNγ somehow “primes” MDMs for inflammasome activation (**Supp Fig. 1**).

**Figure 1.**
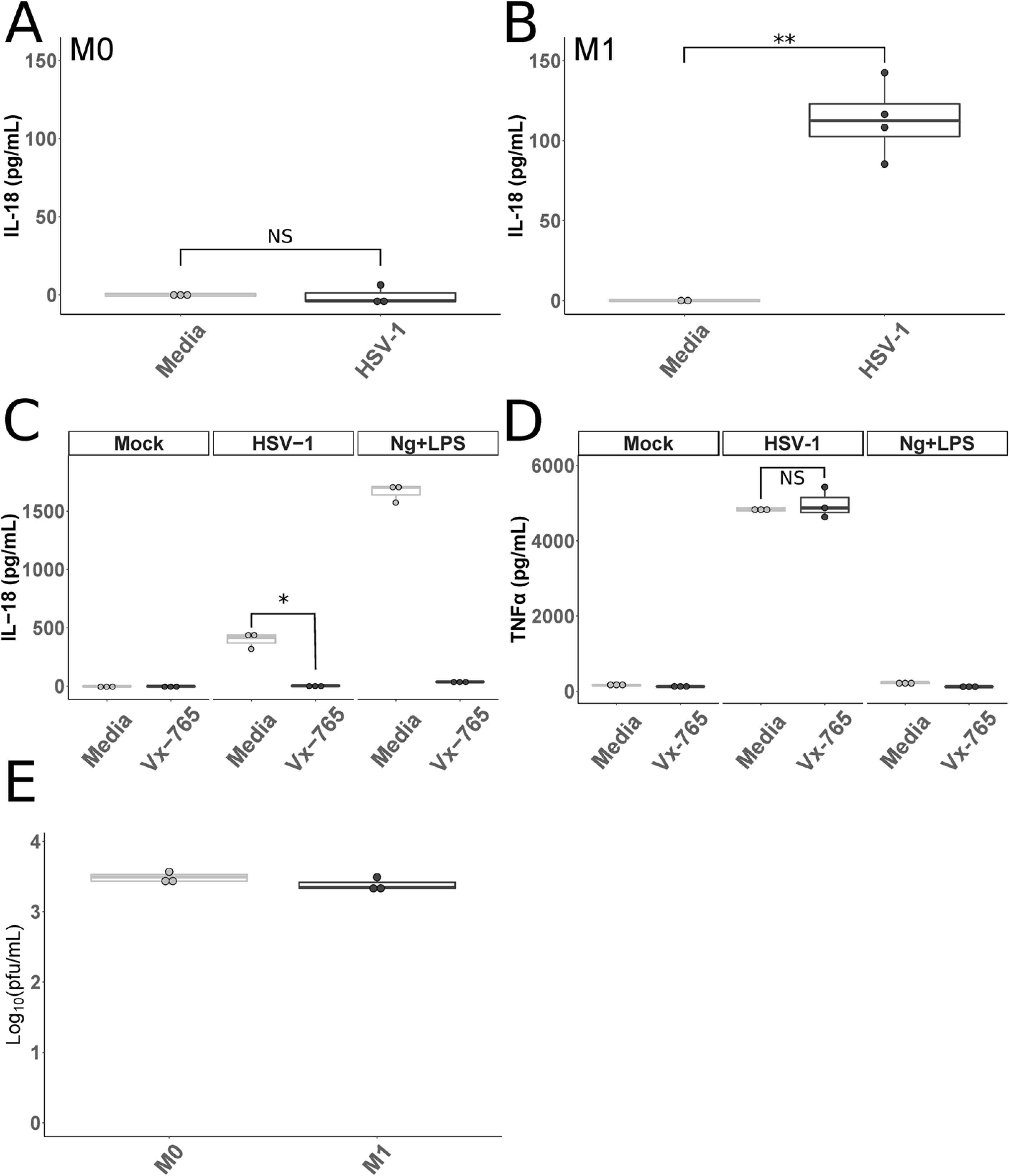
HSV-1 Activates Inflammasomes in Primary Human Macrophages. **A and B.** Primary human MDMs cultured without (M0 **A**) or with IFNγ (M1 **B**) were infected with HSV-1 or mock infected for 24 hours. Cell culture supernatants were collected and assayed for IL-18. **C and D.** Primary human MDMs stimulated with IFNγ were cultured in media alone or media containing 100 ug/mL of VX-765 (Invivogen, San Diego, California) and then inoculated with HSV-1, nigericin and LPS (Ng+LPS), or media as outlined in Materials and Methods for 24 hours. Cell culture supernatants were collected and assayed for IL-18 (**C**) and TNFα (**D**). **E.** MDMs cultured without (M0) or with IFNγ (M1) were infected with HSV-1 for 1 hour followed by citrate wash to inactivate any extracellular virus. Supernatants were collected 24 hours later and PFU were determined via standard plaque assay on Vero cells. Differences between groups indicated by brackets were determined by a student’s t-test. NS, *,**,*** indicate p-values >0.05, <0.05, <0.01, <0.001, respectively.

In order to ensure that this production of IL-18 was dependent on caspase-1, M1 MDMs were treated with VX-765, a caspase-1 specific inhibitor [25–27], prior to either infection with HSV-1 or treatment with nigericin and LPS. IL-18 levels in cell supernatants were reduced to levels not significantly different than background in the presence of VX-765, suggesting that this IL-18 production is due to canonical caspase-1 activation (**Fig. 1c**). To ensure that VX-765 was neither toxic to the cells nor non-specifically interfering with HSV-1 sensing by the macrophages, the same supernatants were tested for TNFα, which is produced by macrophages in response to HSV-1 infection [28]. There was no significant difference in the amount of TNFα produced after HSV-1 infection of macrophages incubated with or without VX-765 (**Fig. 1d**).

To ensure that HSV-1 can replicate in macrophages exposed to IFNγ, M0 MDMs and M1 MDMs were infected with HSV-1, culture supernatants were collected after 24 hours, and plaque forming units (PFU) were determined using a standard plaque assay. HSV-1 was capable of replicating in both resting macrophages and those exposed to IFNγ (**Fig. 1e**). These results demonstrate that IFNγ skewing does not prevent HSV-1 replication in macrophages.

### NLRP3, ASC, and Caspase-1 are required for inflammasome activation in response to HSV-1

To confirm that HSV-1 is capable of activating the inflammasome in a monocyte/macrophage cell line so that dependence on specific inflammasome proteins could be assessed, THP-1 cells were primed with phorbol 12-myristate 13-acetate (PMA) overnight and then infected with HSV-1. IL-18 was measured in supernatants after 24 hours. As previously reported [29], THP-1 cells produced IL-18 after infection with HSV-1 (**Fig. 2a**).

**Figure 2.**
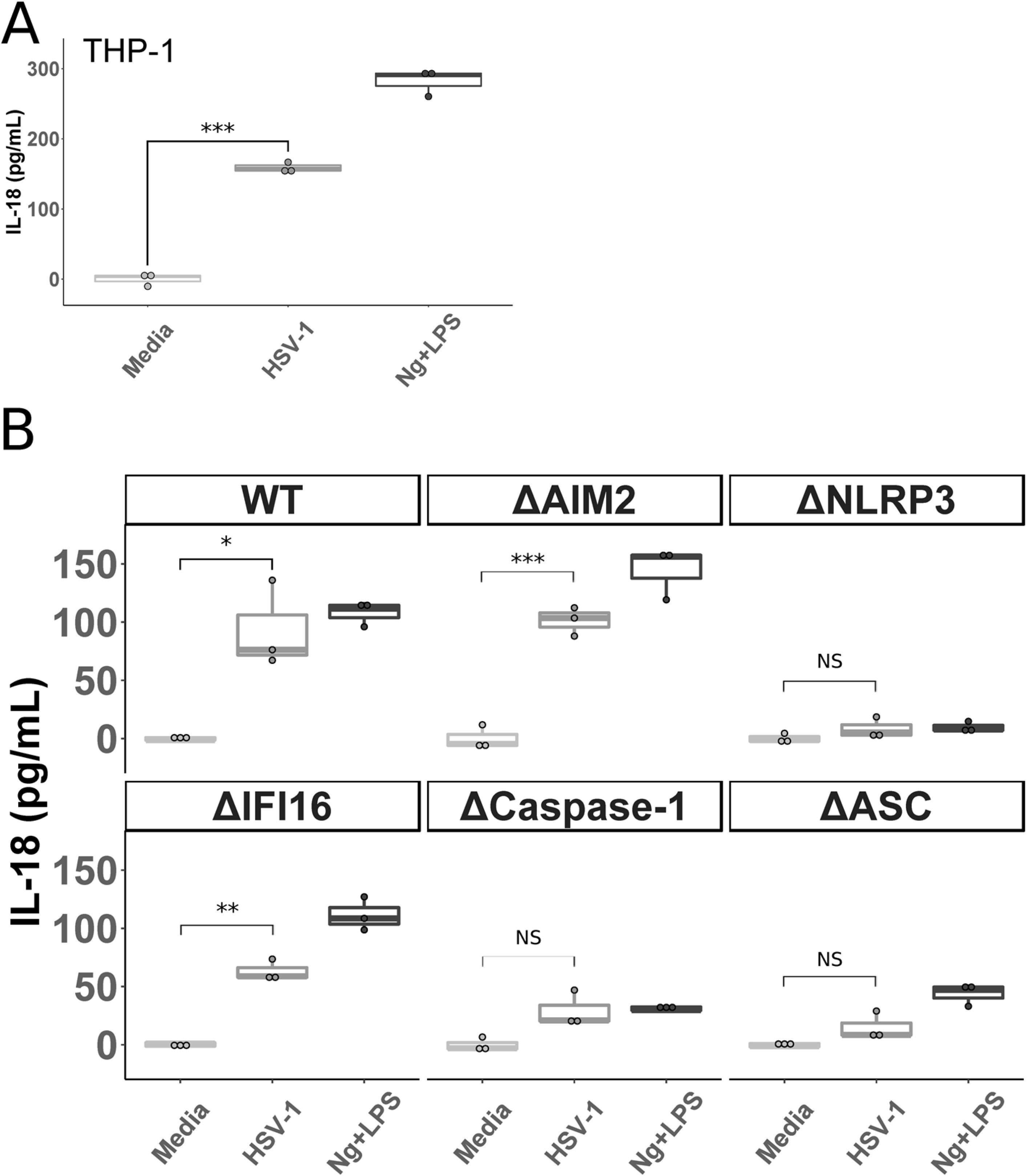
HSV-1 Inflammasome Activation in THP-1 Cells is Dependent on NLRP3, ASC, and Caspase-1. **A.** THP-1 cells were stimulated overnight with PMA (100 ng/mL) and then inoculated with HSV-1, nigericin and LPS (Ng+LPS), or media, as outlined in Materials and Methods, for 24 hours. Cell culture supernatants were collected and IL-18 was measured via ELISA. **B.** THP-1 cells with the indicated gene disrupted via CRISPR-cas9 (Δ) were stimulated overnight with PMA and then inoculated with HSV-1, nigericin and LPS (Ng+LPS), or media for 24 hours before IL-18 was measured in cell supernatants. ΔHUMCYC cells are labeled as “WT.” Differences between groups indicated by brackets were determined by a student’s t-test. NS, *,**,*** indicate p-values >0.05, <0.05, <0.01, <0.001, respectively.

Studies in keratinocytes and human foreskin fibroblasts (HFF) found roles for IFI16, NLRP3, and AIM2 in HSV-1 inflammasome activation [30,31]. Yet, monocytes/macrophages produce the majority of inflammasome related cytokines (IL-18 and IL-1β) in other viral infections and play crucial roles in preventing the most severe manifestations of HSV infection in mouse models [13,32]. Therefore, to determine what inflammasome components are required for HSV-1 induced inflammasome activation in macrophages, we infected THP-1 cells lacking various inflammasome proteins. These cells were constructed using the CRISPR-Cas9 system and previously used to determine the requirements for human cytomegalovirus (HCMV) inflammasome activation in macrophages [33]. The ΔHUMCYC cell-line (WT) was used to control for any off-target effects of the CRISPR-cas9 system. This line was derived from the same THP-1 cells, but targeted a human pseudogene (HUMCYCPS3). While HSV-1 infection of the WT, ΔAIM2, and ΔIFI16 THP-1 cells led to significant IL-18 production, infection of ΔNLRP3, Δcaspase-1, and ΔASC cells resulted in levels of IL-18 that were not significantly different from mock infection (**Fig 2b**). The combination of nigericin with LPS was used as a positive control. As expected, IL-18 concentrations in supernatants from cells lacking NLRP3, ASC, and caspase-1 were not above background after nigericin and LPS exposure. These results indicate that HSV-1 induced inflammasome activation with IL-18 production and release in macrophages is dependent on ASC, caspase-1, and NLRP3, but not on the dsDNA sensors IFI16 or AIM2.

### UV-Irradiated HSV-1 Increases Inflammasome Activation

The HSV-1 tegument protein VP22 blocks activation of the AIM2 inflammasome [21] and; therefore, it is unsurprising that we failed to find a dependence on AIM2. However, it is possible that HSV-1 has evolved multiple mechanisms to alter inflammasome activation. To test this hypothesis, we infected M0 and M1 MDMs with HSV-1 or UV irradiated HSV-1 (HSV-1/UV). Interestingly, HSV-1/UV did lead to IL-18 production in M0 macrophages (**Fig. 3a**). This result suggests that M0 macrophages are capable of inflammasome formation in response to HSV-1, but that a viral factor that is produced during the replication cycle (such as VP22) inhibits this activation. When tested in M1 macrophages, HSV-1/UV led to significantly increased IL-18 production compared to HSV-1 (**Fig 3b**). These data suggest that when macrophages are skewed toward an inflammatory state with IFNγ, a cellular factor is either produced or upregulated that counteracts the inhibitory mechanism(s) of the virus in M0 macrophages. However, replication of the virus does continue to lead to some downregulation of inflammasome activation in IFNγ-primed macrophages because HSV-1/UV led to increased IL-18 production versus HSV-1. One explanation for this phenomenon is that UV-irradiating the virus eliminates sufficient production of VP22 such that AIM2 is able to sense the viral DNA and trigger inflammasome formation in M0s. Whereas the replication competent virus inhibits AIM2 via VP22, the M0s lack additional factor(s) required to trigger inflammasome signaling in response to HSV-1. After skewing with IFNγ, HSV-1 infection leads to inflammasome formation through a non-AIM2 dependent mechanism and HSV-1/UV is able to trigger inflammasome signaling through both AIM2 dependent and non-AIM2 dependent mechanisms.

**Figure 3.**
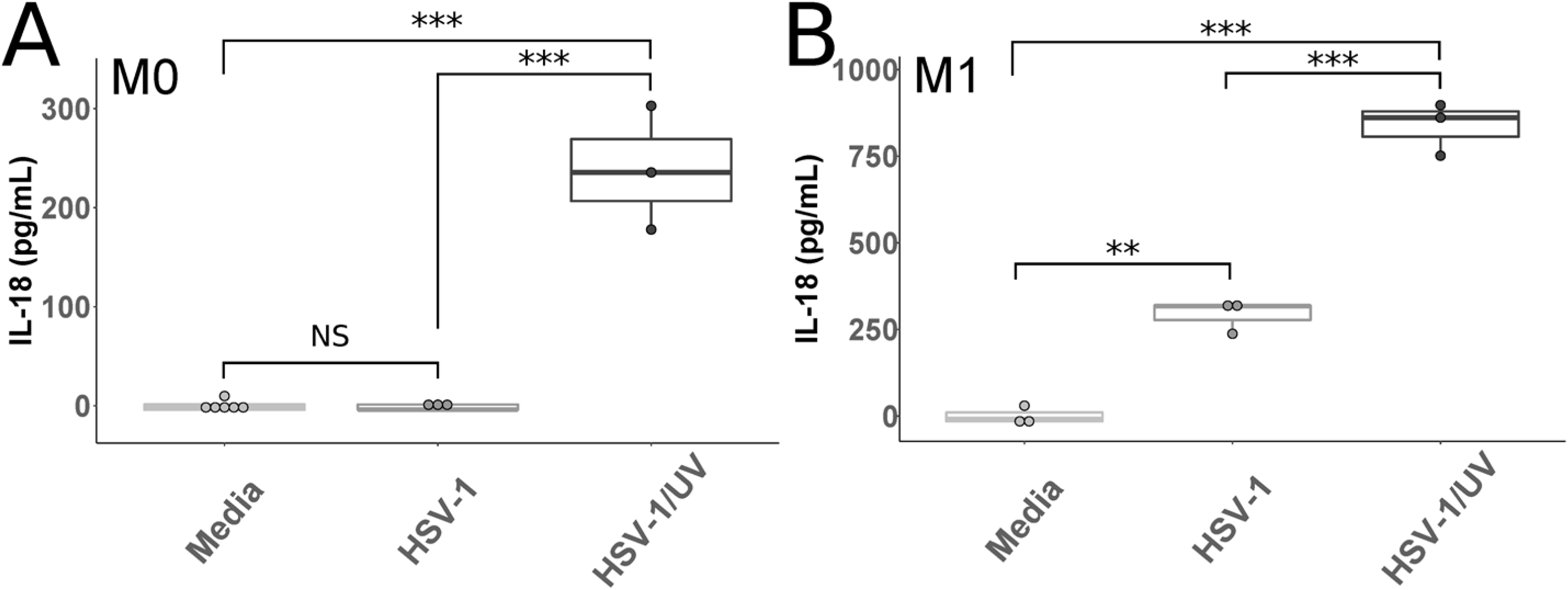
IL-18 in MDMs After Infection with UV-Irradiated HSV-1. **A and B.** Primary human MDMs cultured without (M0 **A**) or with IFNγ (M1 **B**) were infected with HSV-1, UV irradiated HSV-1 (HSV-1/UV), or mock infected for 24 hours. Cell culture supernatants were collected and assayed for IL-18. Differences between indicated conditions within a cell type were determined by a one-way ANOVA with Tukey HSD post-hoc analysis. NS, ****** indicate p-values >0.05, <0.05, <0.01, <0.001 respectively.

To determine if HSV-1 replication in macrophages results in inhibition of any non-AIM2 inflammasome proteins, we tested HSV-1/UV infection of the THP-1 cells lacking AIM2 and NLRP3. In order to more closely replicate the MDM model with IFNγ stimulation, in this experiment the cells were stimulated with PMA and then either infected directly or stimulated with IFNγ for an additional 24hrs and then infected (**Fig 4**). Similar to what was seen in the MDM model, the THP-1 cells that received PMA alone produced only small amounts of IL-18 in response to HSV-1, but significantly more in response to HSV-1/UV. Furthermore, the WT, ΔAIM2, and ΔNLRP3 cells primed with PMA alone all showed a similar increase in IL-18 production in response HSV-1/UV compared to replication competent HSV-1 (**Fig 4a**). Interestingly, after the addition of IFNγ, HSV-1 infection led to significant IL-18 production in the WT and ΔAIM2 cells, with even greater IL-18 produced with exposure to HSV-1/UV. Again, the ΔNLRP3 cells did not produce IL-18 in response to HSV-1, but did produce a modest, but statistically significant, amount of IL-18 after infection with HSV-1/UV (**Fig 4b**). These data confirm our findings in the MDMs that HSV-1 infection of unstimulated macrophages does not lead to inflammasome activation. Further, they support the hypothesis that replication competent HSV-1 is capable of decreasing both AIM2- and NLRP3 dependent inflammasome activation because UV irradiating the virus led to significant increases in IL-18 release in both the ΔAIM2 and ΔNLRP3 lines.

**Figure 4.**
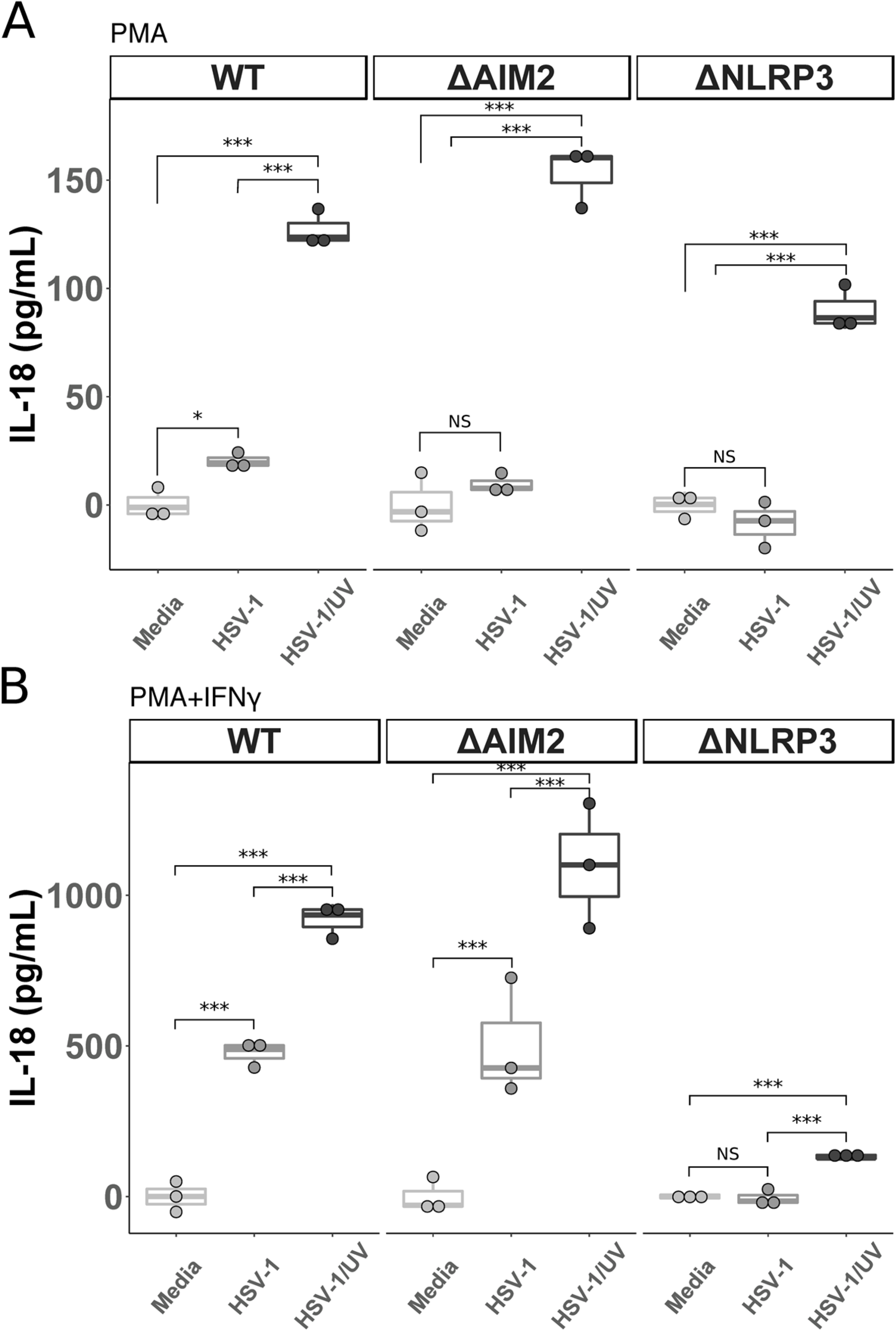
IL-18 in THP-1 Cells After Infection with UV-Irradiated HSV-1. **A.** THP-1 cells were stimulated overnight with PMA (5 ng/mL) and then inoculated with HSV-1, UV irradiated HSV-1 (HSV-1/UV), or mock infected for 24 hours. Cell culture supernatants were collected and IL-18 was measured via ELISA. B. THP-1 cells were stimulated with PMA (5 ng/mL) and then with IFNγ (25 ng/mL) the following day for 24 hours prior to inoculation with HSV-1, UV irradiated HSV-1 (HSV-1/UV), or media alone for 24 hours. Cell culture supernatants were collected and IL-18 was measured via ELISA. ΔHUMCYC cells are labeled as “WT.” Differences between indicated conditions within a cell type were determined by a one-way ANOVA with Tukey HSD post-hoc analysis. NS, *,**,*** indicate p-values >0.05, <0.05, <0.01, <0.001 respectively.

## Discussion

In this study, we demonstrate for the first time that HSV-1 induces IL-18 production and activation of inflammasomes in primary human macrophages stimulated with IFNγ through a caspase-1 dependent process. UV irradiating the virus prior to infection also leads to IL-18 production in unstimulated primary macrophages, but replication competent HSV-1 does not result in IL-18 release without pre-treatment with IFNγ. Furthermore, using THP-1 cell lines, we show that HSV-1 induced inflammasome activation is dependent on NLRP3, ASC, and caspase-1. By comparing HSV-1 and HSV-1/UV in these THP-1 cells, we also provide evidence that HSV-1 is capable of decreasing inflammasome activation through AIM2 and NLRP3 dependent mechanisms. Additionally, our data indicate that a cellular factor is upregulated in macrophages stimulated with IFNγ that allows for activation of the inflammasome after HSV-1 infection. While we do not know what factor is modulated, it is unlikely that it is NLRP3 itself as prior studies have demonstrated no significant increases in NLRP3 expression in macrophages after skewing with IFNγ [23,34]. Finally, activation of the inflammasome in macrophages does not prevent viral replication as measured by plaque assay on macrophage culture supernatants 24 hours after infection with HSV-1.

Although a previous study suggested that HSV-1 infection of primary human macrophages does not lead to inflammasome activation [35], the macrophages in that study were only stimulated with the TLR2 agonist Pam_3_Cys and not IFNγ. Our data in M0-like MDMs showing that HSV-1 infection failed to induce IL-18 secretion are in agreement with this previous study. HSV-1 has been reported to stimulate multiple inflammasome adapter proteins in non-macrophage cell types. In HFFs, HSV-1 was shown to stimulate inflammasome activation through NLRP3 and IFI16 [31] and in keratinocytes it was suggested that HSV-1 activates via NLRP3, IFI16, and AIM2 [30]. However, in our study we found that HSV-1 inflammasome activation in macrophages is dependent on NLRP3 and not IFI16 or AIM2. It is possible that different cell types utilize different inflammasome signaling mechanisms in response to pathogens and that HSV-1 does activate the inflammasome through IFI16 or AIM2 in non-macrophage cells. A recent study in which wild-type THP-1 cells were infected with several strains of HSV-1 showed that more virulent strains of HSV-1 induced more mature IL-18 (measured by western blot) and that multiple inflammasome adapter proteins were upregulated after HSV-1 infection, including NLRP3, NLRP6, NLRP12, and IFI16 [18]. However, it is known that HSV-1 infection leads to upregulation of multiple pro-inflammatory genes and; therefore, increased expression of these inflammasome related proteins does not necessarily indicate that inflammasome signaling is taking place through these adapters [35–37]. Although CMV, a closely related herpesvirus, was recently discovered to activate the inflammasome through AIM2 [33], initial studies on HSV-1 inflammasome activation in macrophages did not find a dependence on AIM2 [20]. This finding was explained by the discovery that VP22 specifically inhibits the interaction between AIM2 and the HSV genome [21]. In agreement with these studies, our current investigation found that AIM2 was not required for HSV-1 to activate the inflammasome in THP-1 cells. Moreover, UV-irradiated HSV-1 led to more IL-18 production, suggesting more robust inflammasome signaling in response to UV-irradiated HSV-1 in both IFNγ stimulated and unstimulated macrophages. UV-irradiated virus is unable to produce de-novo VP22 and therefore the virus is able to activate the inflammasome both through AIM2 and NLRP3. To further support this, while the ΔNLRP3 THP-1 cells did not produce IL-18 in response to HSV-1, they did after UV-irradiating the virus. In this condition, there is insufficient VP22 present to inhibit AIM2 and thus the macrophages are able to sense the HSV-1 genome via AIM2. Interestingly, UV-irradiating HSV-1 prior to infection also led to a robust increase in IL-18 release in the ΔAIM2 THP-1 cell line compared to WT virus. If the VP22-AIM2 interaction is the only mechanism by which HSV-1 is capable of inhibiting inflammasome activation, we would expect there to be no difference in IL-18 production between ΔAIM2 cells infected with replication competent HSV-1 or UV-irradiated HSV-1 because AIM2 is not present. However, we found that UV-irradiating the virus led to an increase in IL-18 in ΔAIM2 THP-1 cells, suggesting that the virus has evolved other mechanisms to inhibit inflammasome activation in macrophages that are not AIM2 dependent. Having multiple mechanisms of evastion highlights the importance of the role of inflammasome activation in macrophages to control HSV-1.

The primary limitation of our study is that it was restricted to primary macrophages and macrophage-like cell lines. As discussed, our data support that HSV-1 is capable of activating more than one inflammasome signaling adapter and the signaling pathway may differ depending on the cell type studied. Therefore, we cannot draw conclusions regarding the interaction between HSV-1 and IFI16 or other inflammasome related proteins in all cell types that the virus is capable of infecting. However, macrophages are a crucial cell type in inflammasome activation and HSV-1 control in murine models, prompting our focus on this cell type. Additionally, the present studies were centered on human cells and cell lines and we did not investigate these inflammasome proteins in other species or whole animal models. A prior study in mice showed that HSV-1 causes more severe keratitis after corneal infection in NLRP3 KO mice compared to WT [38]. This suggests that regulation of this pathway is central to the delicate balance between viral control and excessive tissue damage.

In summary, we have demonstrated that HSV-1 infection leads to production of IL-18 through canonical caspase-1 inflammasome activation in primary human macrophages. This process is dependent on the inflammasome proteins NLRP3, ASC, and caspase-1. Furthermore, our data demonstrate that HSV-1 replication partially inhibits NLRP3 dependent inflammasome activation in human cells.

## Materials and Methods

### Cells and Viruses

HSV-1 strain KOS was the generous gift of Richard Longnecker (Northwestern University). Virus was propagated in Vero cells (also a gift from Richard Longnecker, Northwestern University) cultured in Dulbecco’s modification of Eagle medium with 1% fetal bovine serum (DMEV) as previously described [39,40]. Standard plaque titrations to determine viral titers were performed on confluent monolayers of Vero cells. For UV activation the inoculum was dispensed in a sterile basin in a biosafety cabinet with a UV lamp source (Sankyo Denki G30T8) and irradiated for 30 minutes. Virus inactivation was confirmed by standard plaque assay. Titer decrease of ≥ 10^6^ PFU/mL was considered successful. Cells were incubated at 37°C and 5% CO_2_ unless otherwise stated. Vero cells were the generous gift of Richard Longnecker (Northwestern University) and were maintained in Dulbecco’s modification of Eagle medium with 10% fetal bovine serum and 1% penicillin/streptomycin (DME). PBMCs were isolated by Ficoll-Hypaque gradient centrifugation. Primary monocytes were magnetically sorted by negative isolation per the manufacturer’s specifications (Miltenyi Biotec, Somerville, Massachusetts) and cultured in RPMI 1640 (Invitrogen, Waltham, Massachusetts) with 10% heat-inactivated fetal bovine serum, 1% Penicillin/Streptomycin, L-glutamine (2mM) and 50 ng/mL of recombinant human M-CSF (R&D Systems, Minneapolis, Minnesota) for 6 to 7 days to differentiate them to macrophages [41]. Adherent macrophages were washed with sterile PBS and then incubated with the non-enzymatic cell disassociation media, CellStripper (Corning, Tewksbury, Massachusetts), for 30 minutes at 37°C and 5% CO_2_ followed by counting, centrifugation at 400g for 5 min, and plating at a density of 3×10^5^ cells/well in a sterile U-bottom 96-well plate (unless otherwise stated). For M1 differentiation, macrophages were cultured overnight in RPMI 1640 with 10% heat-inactivated fetal bovine serum, 1% Penicillin/Streptomycin, L-glutamine (2mM), and IFNγ (25ng/mL) (Peprotech, Rocky Hill, New Jersey) [34]. M0 macrophages were cultured in the same base media, but without IFNγ. The generation of the THP-1 cells was well described previously [33]. THP-1 cells were maintained in RPMI 1640 media, 10% heat-inactivated fetal bovine serum, 1 % MEM nonessential amino acids, 1% Penicillin/Streptomycin, sodium pyruvate, and L-glutamine (2mM) at a density of 5×10^5^ – 2×10^6^ cells/mL. To differentiate into macrophages THP-1 cells were plated at a density of 3×10^5^ cells/well in a sterile U-bottom 96-well plate and stimulated overnight in RPMI 1640 media, 2% heat-inactivated fetal bovine serum, 1% Penicillin/Streptomycin, L-glutamine (2mM), and phorbol 12-myristate 13-acetate (PMA) 100 ng/mL (unless otherwise stated).

### Infections and Inflammasome Activation

Unless otherwise stated, all infections were carried out at a multiplicity of infection (MOI) of 10. For both HSV-1 infection and nigericin stimulation, media was gently aspirated from the cell culture wells containing the indicated cells and replaced with RPMI 1640 media containing 2% heat-inactivated fetal bovine serum (R2) and either HSV-1 (at a MOI of 10), nigericin (5uM) (MilliporeSigma, Burlington, Massachusetts) and LPS (1ug/mL), or no additional reagents (mock/media control). Twenty-four hours later supernatant was removed and used for downstream assays. To measure viral progeny produced in macrophages (**Fig 4**), macrophages were plated in sterile 12-well culture dishes at a density if 5×10^5^ cells/well. Media was aspirated and replaced with HSV-1 strain KOS in R2 and incubated at 37°C for 1 hr. The inoculum was aspirated, cells were washed with sterile phosphate-buffered saline (PBS), washed with a citrate solution (pH 3) to inactivate any viral particles that had not entered, and fresh warm R2 was added back to the cells. 24 hours later cell culture media was harvested and PFU were determined as described above.

### IL-18 and TNFa Measurements

Human IL-18 and TNFα were measured with the human IL-18 ELISA Kit (MBL, Woburn, Massachusetts) and human TNFα ELISA Kit (ThermoFisher, Waltham, Massachusetts) according to the manufacturers’ instructions using cell culture supernatant at a 1:5 dilution. The lower limit of detection was 12 pg/mL for IL-18 and 7.8 pg/mL for TNFα. Data were acquired on a SpectraMax M2. Results were analyzed using R. Unless otherwise stated, all measurements were normalized to the average of the media control for each experiment.

### Ethics Statement

For experiments involving primary human macrophages, deidentified human blood Leuko Paks were obtained from the Anne Arundel Medical Blood Donor Center (Anne Arundel, Maryland, USA).

## Acknowledgments

We thank members of the Viral Hepatitis Center at Johns Hopkins for advice and discussion particularly Michael Chattergoon, Laura Cohen, Kim Rousseau, and Katie Cascino. We thank Richard Longnecker and Nan Susmarski at Northwestern University for cells and viruses. Financial Support: This work was supported by the National Institute of Allergy and Infectious Diseases U19AI088791 and R01AI108403. AHK was supported by the National Institute of Health T32 AI007291-27. AF was supported, in part, by grant D18HP29037 from the U.S. Health Resources and Services Administration, Bureau of Health Workforce, Health Careers Opportunity Program. The content is solely the responsibility of the authors and does not necessarily represent the official views of the National Institutes of Health.

## Author Contributions

AHK: Conceptualization, Investigation, Formal Analysis, Visualization, Writing – Original Draft Preparation

AF: Conceptualization, Investigation, Writing – Review and Editing

GM: Conceptualization, Investigation

SB: Resources, Investigation

VRD: Resources, Writing – Review and Editing

ALC: Conceptualization, Funding Acquisition, Methodology, Resources, Supervision, Writing – Review and Editing

## Supporting Information Legends

**Supplemental Figure 1.**
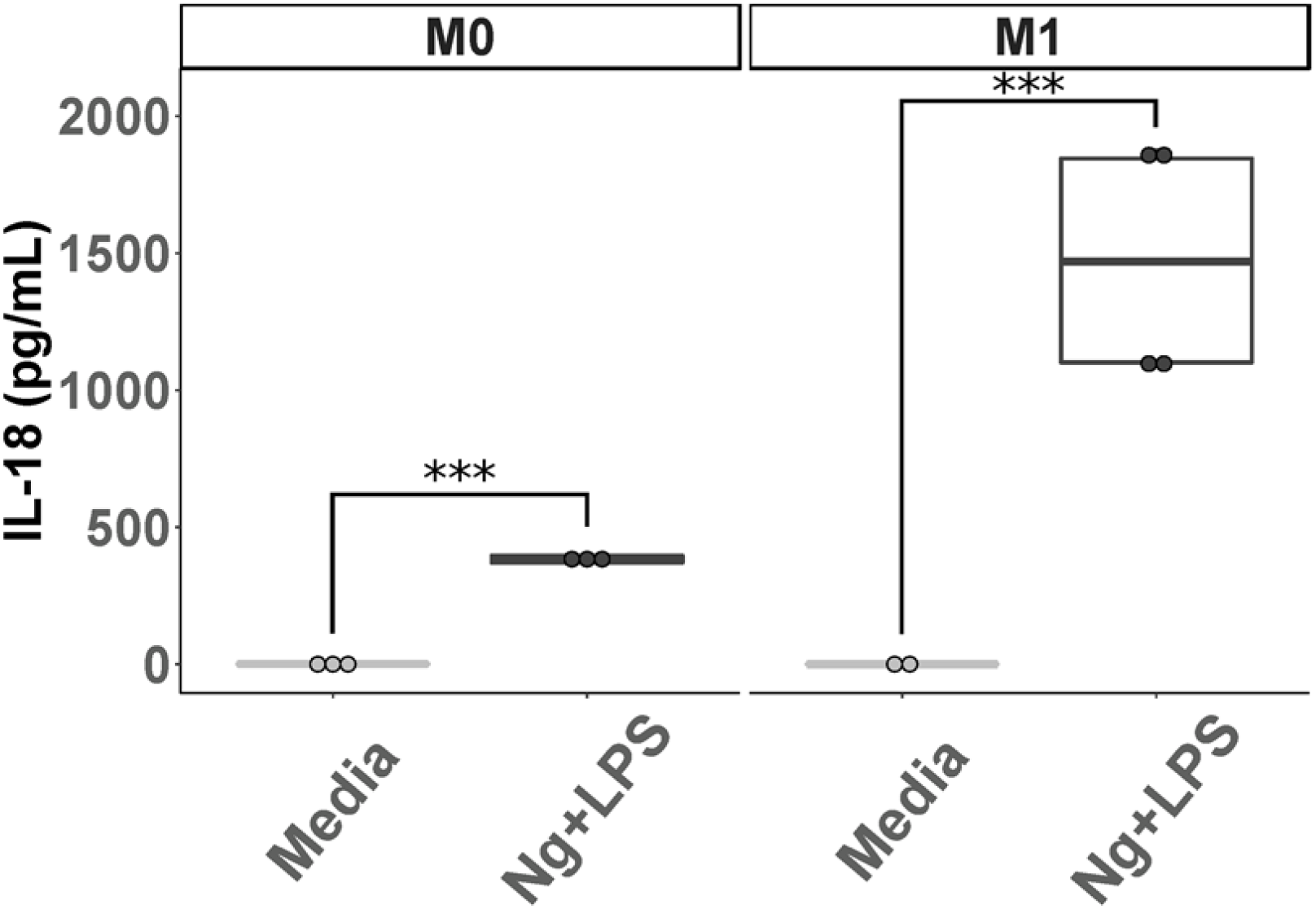
Nigericin Activates Inflammasomes in Primary Human Macrophages. Primary human MDMs cultured without (M0 left panel) or with IFNγ (M1 right panel) were incubated with nigericin and LPS (Ng+LPS), or media, as outlined in Materials and Methods, for 24 hours. Cell culture supernatants were collected and assayed for IL-18. Differences between groups indicated by brackets were determined by a student’s t-test. NS, *,**,*** indicate p-values >0.05, <0.05, <0.01, <0.001, respectively.

## References

1. Brubaker SW, Bonham KS, Zanoni I, Kagan JC. Innate Immune Pattern Recognition: A Cell Biological Perspective. http://dxdoiorg/101146/annurev-immunol-032414-112240. Annual Reviews; 2015;33: 257–290. doi:10.1146/annurev-immunol-032414-112240

2. Frank D, Vince JE. Pyroptosis versus necroptosis: similarities, differences, and crosstalk. Cell Death Differ. Nature Publishing Group; 2019;26: 99–114. doi:10.1038/s41418-018-0212-6

3. Mantovani A, Dinarello CA, Molgora M, Garlanda C. Interleukin-1 and Related Cytokines in the Regulation of Inflammation and Immunity. Immunity. Elsevier Inc; 2019;50: 778–795. doi:10.1016/j.immuni.2019.03.012

4. Costa LS, Outlioua A, Anginot A, Akarid K, Arnoult D. RNA viruses promote activation of the NLRP3 inflammasome through cytopathogenic effect-induced potassium efflux. Cell Death and Disease. Springer US; 2019;: 1–15. doi:10.1038/s41419-019-1579-0

5. Chen W, Xu Y, Li H, Tao W, Xiang Y, Huang B, et al. HCV Genomic RNA Activates the NLRP3 Inflammasome in Human Myeloid Cells. Sutterwala FS, editor. PLoS ONE. Public Library of Science; 2014;9: e84953–10. doi:10.1371/journal.pone.0084953

6. de Castro-Jorge LA, de Carvalho RVH, Klein TM, Hiroki CH, Lopes AH, Guimarães RM, et al. The NLRP3 inflammasome is involved with the pathogenesis of Mayaro virus. Randall G, editor. PLoS Pathog. Public Library of Science; 2019;15: e1007934–27. doi:10.1371/journal.ppat.1007934

7. Kuriakose T, Kanneganti T-D. Regulation and functions of NLRP3 inflammasome during influenza virus infection. Mol Immunol. 2017;86: 56–64. doi:10.1016/j.molimm.2017.01.023

8. Man SM, Karki R, Kanneganti T-D. Molecular mechanisms and functions of pyroptosis, inflammatory caspases and inflammasomes in infectious diseases. Immunol Rev. 2017;277: 61–75. doi:10.1111/imr.12534

9. Voet S, Srinivasan S, Lamkanfi M, van Loo G. Inflammasomes in neuroinflammatory and neurodegenerative diseases. EMBO Mol Med. 2019. doi:10.15252/emmm.201810248

10. Chen I-Y, Ichinohe T. Response of host inflammasomes to viral infection. Trends Microbiol. 2015;23: 55–63. doi:10.1016/j.tim.2014.09.007

11. Broz P, Dixit VM. Inflammasomes: mechanism of assembly, regulation and signalling. Nat Rev Immunol. 2016;16: 407–420. doi:10.1038/nri.2016.58

12. Gaidt MM, Ebert TS, Chauhan D, Schmidt T, Schmid-Burgk JL, Rapino F, et al. Human Monocytes Engage an Alternative Inflammasome Pathway. Immunity. Elsevier Inc; 2016;44: 833–846. doi:10.1016/j.immuni.2016.01.012

13. Chattergoon MA, Latanich R, Quinn J, Winter ME, Buckheit RW, Blankson JN, et al. HIV and HCV Activate the Inflammasome in Monocytes and Macrophages via Endosomal Toll-Like Receptors without Induction of Type 1 Interferon. Paludan SR, editor. PLoS Pathog. Public Library of Science; 2014;10: e1004082–12. doi:10.1371/journal.ppat.1004082

14. Fields BN, Knipe DM, Howley PM, Technologies IO. Fields virology. 6 ed. Philadelphia: Wolters Kluwer/Lippincott Williams & Wilkins Health; 2013.

15. Horan KA, Hansen K, Jakobsen MR, Holm CK, Søby S, Unterholzner L, et al. Proteasomal degradation of herpes simplex virus capsids in macrophages releases DNA to the cytosol for recognition by DNA sensors. The Journal of Immunology. 2013;190: 2311–2319. doi:10.4049/jimmunol.1202749

16. Ghiasi H, Wechsler SL, Kaiwar R, Nesburn AB, Hofman FM. Local expression of tumor necrosis factor alpha and interleukin-2 correlates with protection against corneal scarring after ocular challenge of vaccinated mice with herpes simplex virus type 1. Journal of Virology. American Society for Microbiology Journals; 1995;69: 334–340.

17. Lee AJ, Chen B, Chew MV, Barra NG, Shenouda MM, Nham T, et al. Inflammatory monocytes require type I interferon receptor signaling to activate NK cells via IL-18 during a mucosal viral infection. J Exp Med. 2017;214: 1153–1167. doi:10.1084/jem.20160880

18. Coulon P-G, Dhanushkodi N, Prakash S, Srivastava R, Roy S, Alomari NI, et al. NLRP3, NLRP12, and IFI16 Inflammasomes Induction and Caspase-1 Activation Triggered by Virulent HSV-1 Strains Are Associated With Severe Corneal Inflammatory Herpetic Disease. Front Immunol. Frontiers; 2019;10: 38–19. doi:10.3389/fimmu.2019.01631

19. Sergerie Y, Rivest S, Boivin G. Tumor Necrosis Factor–α and Interleukin-1β Play a Critical Role in the Resistance against Lethal Herpes Simplex Virus Encephalitis. J Infect Dis. 2007;196: 853–860. doi:10.1086/520094

20. Rathinam VAK, Jiang Z, Waggoner SN, Sharma S, Cole LE, Waggoner L, et al. The AIM2 inflammasome is essential for host defense against cytosolic bacteria and DNA viruses. Nat Immunol. Nature Publishing Group; 2010;11: 395. doi:10.1038/ni.1864

21. Maruzuru Y, Ichinohe T, Sato R, Miyake K, Okano T, Suzuki T, et al. Herpes Simplex Virus 1 VP22 Inhibits AIM2-Dependent Inflammasome Activation to Enable Efficient Viral Replication. Cell Host Microbe. 2018;23: 254–265.e7. doi:10.1016/j.chom.2017.12.014

22. Martinez FO, Gordon S, Locati M, Mantovani A. Transcriptional Profiling of the Human Monocyte-to-Macrophage Differentiation and Polarization: New Molecules and Patterns of Gene Expression. J Immunol. American Association of Immunologists; 2006;177: 7303–7311. doi:10.4049/jimmunol.177.10.7303

23. Beyer M, Mallmann MR, Xue J, Staratschek-Jox A, Vorholt D, Krebs W, et al. High-Resolution Transcriptome of Human Macrophages. Zirlik A, editor. PLoS ONE. Public Library of Science; 2012;7: e45466–16. doi:10.1371/journal.pone.0045466

24. He Y, Hara H, Nuñez G. Mechanism and Regulation of NLRP3 Inflammasome Activation. Trends Biochem Sci. 2016;41: 1012–1021. doi:10.1016/j.tibs.2016.09.002

25. Labzin LI, Bottermann M, Rodriguez Silvestre P, Foss S, Andersen JT, Vaysburd M, et al. Antibody and DNA sensing pathways converge to activate the inflammasome during primary human macrophage infection. The EMBO Journal. John Wiley & Sons, Ltd; 2019;6: 37457–16. doi:10.15252/embj.2018101365

26. Doitsh G, Galloway NLK, Geng X, Yang Z, Monroe KM, Zepeda O, et al. Cell death by pyroptosis drives CD4 T-cell depletion in HIV-1 infection. Nature. Nature Publishing Group; 2014;505: 509–514. doi:10.1038/nature12940

27. Tyrkalska SD, Pérez-Oliva AB, Rodríguez-Ruiz L, Martínez-Morcillo FJ, Alcaraz-Pérez F, Martínez-Navarro FJ, et al. Inflammasome Regulates Hematopoiesis through Cleavage of the Master Erythroid Transcription Factor GATA1. Immunity. Elsevier Inc; 2019;51: 50–63.e5. doi:10.1016/j.immuni.2019.05.005

28. Melchjorsen J, Rintahaka J, Søby S, Horan KA, Poltajainen A, Østergaard L, et al. Early innate recognition of herpes simplex virus in human primary macrophages is mediated via the MDA5/MAVS-dependent and MDA5/MAVS/RNA polymerase III-independent pathways. Journal of Virology. American Society for Microbiology Journals; 2010;84: 11350–11358. doi:10.1128/JVI.01106-10

29. Muruve DA, Pétrilli V, Zaiss AK, White LR, Clark SA, Ross PJ, et al. The inflammasome recognizes cytosolic microbial and host DNA and triggers an innate immune response. Nature. Nature Publishing Group; 2008;452: 103–107. doi:10.1038/nature06664

30. Strittmatter GE, Sand J, Sauter M, Seyffert M, Steigerwald R, Fraefel C, et al. IFN-γ Primes Keratinocytes for HSV-1-Induced Inflammasome Activation. J Invest Dermatol. 2016;136: 610–620. doi:10.1016/j.jid.2015.12.022

31. Johnson KE, Chikoti L, Chandran B. Herpes Simplex Virus 1 Infection Induces Activation and Subsequent Inhibition of the IFI16 and NLRP3 Inflammasomes. Journal of Virology. 2013;87: 5005–5018. doi:10.1128/JVI.00082-13

32. Lupfer C, Malik A, Kanneganti T-D. Inflammasome control of viral infection. Current Opinion in Virology. 2015;12: 38–46. doi:10.1016/j.coviro.2015.02.007

33. Botto S, Abraham J, Mizuno N, Pryke K, Gall B, Landais I, et al. Human Cytomegalovirus Immediate Early 86-kDa Protein Blocks Transcription and Induces Degradation of the Immature Interleukin-1β Protein during Virion-Mediated Activation of the AIM2 Inflammasome. mBio. 2019;10. doi:10.1128/mBio.02510-18

34. Awad F, Assrawi E, Jumeau C, Georgin-Lavialle S, Cobret L, Duquesnoy P, et al. Impact of human monocyte and macrophage polarization on NLR expression and NLRP3 inflammasome activation. Lai H-C, editor. PLoS ONE. 2017;12: e0175336–18. doi:10.1371/journal.pone.0175336

35. Miettinen JJ, Matikainen S, Nyman TA. Global secretome characterization of herpes simplex virus 1-infected human primary macrophages. Journal of Virology. 2012;86: 12770–12778. doi:10.1128/JVI.01545-12

36. Ellermann-Eriksen S. Macrophages and cytokines in the early defence against herpes simplex virus. Virol J. 2005;2: 59. doi:10.1186/1743-422X-2-59

37. Tognarelli EI, Palomino TF, Corrales N, Bueno SM, Kalergis AM, González PA. Herpes Simplex Virus Evasion of Early Host Antiviral Responses. Front Cell Infect Microbiol. Frontiers; 2019;9: 8998–24. doi:10.3389/fcimb.2019.00127

38. Giménez F, Bhela S, Dogra P, Harvey L, Varanasi SK, Jaggi U, et al. The inflammasome NLRP3 plays a protective role against a viral immunopathological lesion. J Leukoc Biol. 2016;99: 647–657. doi:10.1189/jlb.3HI0715-321R

39. Karaba AH, Kopp SJ, Longnecker R. Herpesvirus entry mediator and nectin-1 mediate herpes simplex virus 1 infection of the murine cornea. Journal of Virology. 2011;85: 10041–10047. doi:10.1128/JVI.05445-11

40. Karaba AH, Kopp SJ, Longnecker R. Herpesvirus entry mediator is a serotype specific determinant of pathogenesis in ocular herpes. Proc Natl Acad Sci USA. 2012. doi:10.1073/pnas.1216967109

41. Hashimoto S, Yamada M, Motoyoshi K, Akagawa KS. Enhancement of macrophage colony-stimulating factor-induced growth and differentiation of human monocytes by interleukin-10. Blood. 1997;89: 315–321.

